# Unleashing floret fertility by a mutated homeobox gene improved grain yield during wheat evolution under domestication

**DOI:** 10.1101/434985

**Authors:** Shun Sakuma, Guy Golan, Zifeng Guo, Taiichi Ogawa, Akemi Tagiri, Kazuhiko Sugimoto, Nadine Bernhardt, Jonathan Brassac, Martin Mascher, Goetz Hensel, Shizen Ohnishi, Hironobu Jinno, Yoko Yamashita, Idan Ayalon, Zvi Peleg, Thorsten Schnurbusch, Takao Komatsuda

## Abstract

Floret fertility is a key trait to determine the number of grains per inflorescence in cereals. During wheat (*Triticum* sp.) evolution, floret fertility has been increased and current bread wheat (*T. aestivum* L.) produces three to five grains per spikelet; however, little is known about the genetic basis controlling floret fertility. Here we identify the quantitative trait locus *Grain Number Increase 1 (GNI1)*, encoding a homeodomain leucine zipper class I (HD-Zip I) transcription factor. *GNI1* evolved in the Triticeae through gene duplication and functionalization. *GNI1* was predominantly expressed in the most apical floret primordia and parts of the rachilla, suggesting that GNI1 inhibits rachilla growth and development. *GNI1* expression decreased during wheat evolution, and as a consequence, more fertile florets and grains per spikelet are being produced. Genetic analysis revealed that the reduced-function allele of *GNI1-A* contributes to increase the number of fertile florets per spikelet. The knockdown of *GNI1* in transgenic hexaploid wheat improved fertile floret and grain number. Furthermore, wheat plants carrying the impaired allele increased grain yield under field conditions. Our findings illuminate that gene duplication and functionalization generated evolutionary novelty for floret fertility (i.e. reducing floral numbers) while the mutations towards increased grain production were under selection during wheat evolution under domestication.

**Significance Statement:** Grain number is a fundamental trait for cereal grain yield; but its underlying genetic basis is mainly unknown in wheat. Here we show for the first time a direct link between increased floret fertility, higher grain number per spike and higher plot-yields of wheat in the field. We have identified *GNI1* gene encoding an HD-Zip I transcription factor responsible for increased floret fertility. The wild type allele imposes an inhibitory role specifically during rachilla development, indicating that expression of this protein actively shuts-down grain yield potential; whereas, the reduced-function allele enables more florets and grains to be produced. *GNI1* evolved through gene duplication in Triticeae and its mutations were under parallel human selection during wheat and barley evolution under domestication.

## Introduction

The tribe Triticeae within the Pooideae subfamily of the Poaceae family consists of approximately 30 genera with 360 species including economically important cereal crops such as bread wheat (*Triticum aestivum* L.), durum wheat [*T. turgidum* ssp. *durum* (Desf.) MacKey], barley (*Hordeum vulgare* L.) and rye (*Secale cereale* L.) (1). Triticeae plants produce an unbranched inflorescence called spike. While most species develop a single spikelet on each rachis node like wheat; others can produce two or more spikelets on each rachis node (2). The wheat spike is consisting of several spikelets with a terminal spikelet at the apex while each spikelet generates an indeterminate number of florets attached to a secondary axis, called rachilla (3, 4). Final grain number per spikelet is determined by the fertility of each floret within a spikelet (5, 6). At the white anther stage, a wheat spikelet normally produces up to ten to 12 floret primordia (**Fig. 1A**); however, during development more than ~70% of the florets will undergo degeneration. This phenomenon is known as the floret abortion (6, 7). Despite its prominent importance for grain number determination and the potential for grain yield improvement, little is known about the genetic basis of floret fertility in wheat.

**Fig. 1.**
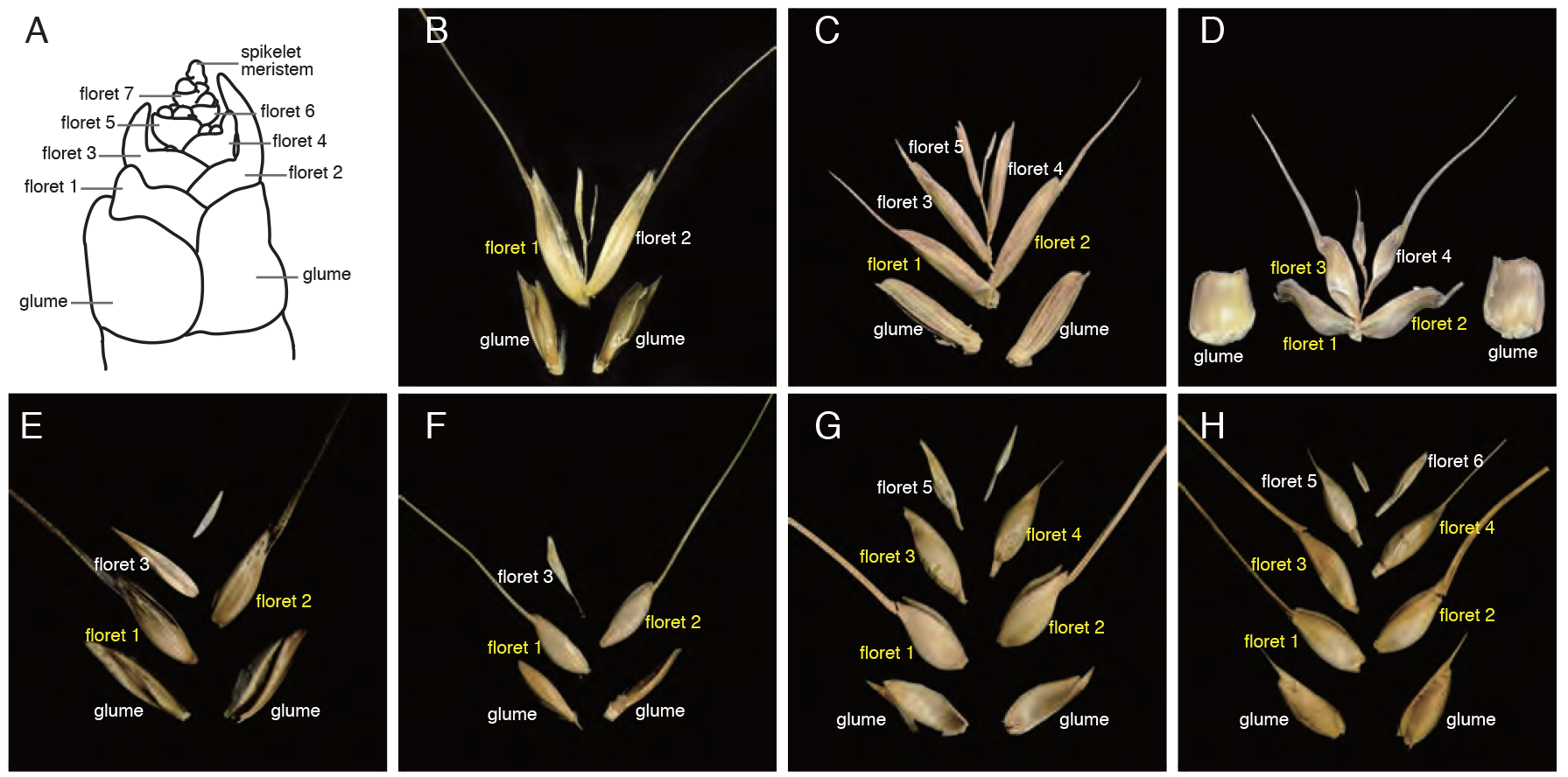
Structure of wheat spikelet. (*A*) Schematic model of wheat spikelet at white anther stage (35). (*B*–*D*) Diploid progenitors of bread wheat. (*B*) *Triticum urartu*. (*C*) *Aegilops speltoides*. (*D*) *Ae. tauschii*. (*E*–-*G*) Tetraploid wheat. (*E*) Wild emmer wheat; *T. turgidum* ssp. *dicoccoides*. (*F*) Domesticated emmer wheat; *T. turgidum* ssp. *diccocum*. (*G*) Durum wheat; *T. turgidum* ssp. *durum*. (*H*) Hexaploid bread wheat; *T. aestivum*. Yellow-labelled florets indicate fertile florets producing grain.

Two polyploidization events produced allohexaploid bread wheat consisting of three subgenomes (B, A, and D)(8, 9). The wild emmer [*T. turgidum* ssp. *dicoccoides* (Körn.) Thell.; 2n=28, BBAA] allotetraploid genome arose about 0.36 to 0.5 million years ago, presumably through a single event of hybridization and spontaneous chromosome doubling. The A-subgenome originated from *T. urartu* (2n=14, AA-genome) and the B-subgenome from *Aegilops speltoides* (2n=14, SS-genome) or an extinct closely-related species. Domesticated emmer wheat (*T. turgidum* ssp. *dicoccum* Schrank) was derived from wild emmer and led to durum wheat, which is most widely cultivated tetraploid wheat. A second polyploidization event ~7,000 yeas ago between domesticated tetraploid wheat and *Ae. tauschii* (2n=14, DD-genome) produced the ancestral allohexaploid wheat (*T. aestivum*; 2n=42, BBAADD). As the ploidy level increases, more floret/grain number per spikelet was produced in *Triticum* species: one or two grains per spikelet in diploid (*T. urartu* and *T. monococcum*), two or three in tetraploid, and more than three in hexaploid wheat (**Fig. 1B-H**) (10).

Genetic diversity of grass inflorescences determines its reproduction system, and hence, number of branches, flowers and grains (11). The grass inflorescences are classified as a “raceme” having one central monopodial axis, a “panicle” with primary and secondary branches, or a “spike” without pedicels (flower stalk). Following cereals domestication, inflorescence architecture has been improved as more reproductive branches, called spikelets, produced more grains (12). Since inflorescence architecture has been a target for selection, a better understanding of the genes that regulate spikelet development could help increase cereal grain yield. The regulation of floret number per spikelet is a main determinant of spikelet architecture. One traditional classification of ‘spikelet’ is whether it contains a determinate or indeterminate number of florets. For example, the spikelet of rice (*Oryza sativa* L.), barley, sorghum (*Sorghum bicolor* L.) and maize (*Zea mays* L.) are determinate, producing only one floret in rice and barley and two florets per spikelet in sorghum and maize. On the other hand, an indeterminate number of florets per spikelet are produced in wheat and oat (*Avena sativa* L.). Interestingly, there are sterile florets in common independent of the determinacy of floret number; e.g. two lateral florets in two-rowed barley, a lower floret in maize and sorghum, or several apical florets in wheat and oat.

Recent studies have suggested that variations in grain number per spike had a larger effect on wheat grain yield than variations in grain size (13, 14). Quantitative trait loci (QTLs) affecting grain number per spike in wheat have been mapped; but their underlying gene(s) are yet to be discovered (15–18). Genome-wide association analyses among European winter wheats revealed a QTL on chromosome 2AL conferring enhanced grain number per spikelet (19); however the responsible gene has not been elucidated. In this study, we investigated natural variations for grain number per spikelet in polyploid wheats and their wild relatives and identify the underlying gene for improved floret fertility and enhanced grain number. We also explored the evolutionary trajectory of floret fertility during wheat evolution.

## Results

### QTL cloning for grain number per spikelet

To understand the genetic regulation in the number of fertile florets per spikelet, a population of recombinant inbred substitution lines derived from a cross between durum wheat cv. Langdon (LDN) and the substitution line DIC-2A(20) was investigated. The DIC-2A contained the chromosome 2A from the wild emmer wheat accession ISR-A in the LDN genetic background and produced two grains per spikelet on average while LDN has 2.4 on average (**Fig. 2A**). Increased grain number per spikelet in LDN was mainly due to increased grains in basal and central spike parts (**Fig. 2B**). QTL analysis detected a single major QTL with a log_10_ odds (LOD) score of 18.71 on chromosome 2AL, explaining 61% of the phenotypic variance (**Fig. 2C**). To further limit the region, backcrossed recombinant lines were developed and the QTL was mapped as a simple Mendelian locus, named *Grain Number Increase 1*, *GNI1-A* (**Fig. 2D**). Fine mapping delimited the *GNI1-A* locus within a 5.4 Mbp interval containing 26 putative genes, including one that encoded an HD-Zip I transcription factor protein, which is the closest wheat homolog of barley *Six-rowed spike 1* gene *vrs1* (21)(***SI Appendix*, Table S1**). Comparison of the parental sequences revealed a single amino acid substitution (N105Y: 105 asparagine to tyrosine) in the highly conserved homeodomain (**Fig. 2E**). The recombinant plants carrying the LDN allele (105Y) showed a significantly higher grain number per spikelet than the plants carrying the DIC-2A allele (105N) mainly due to an additive effect of the LDN allele (**Fig. 2F**). Interestingly, the mutation in LDN was identical to the one already found in barley six-rowed spike mutant *Int-d.41* allelic at the *vrs1* locus (21), strongly suggesting that the HD-Zip I protein encoded by LDN allele most likely lost or reduced its function.

**Fig. 2.**
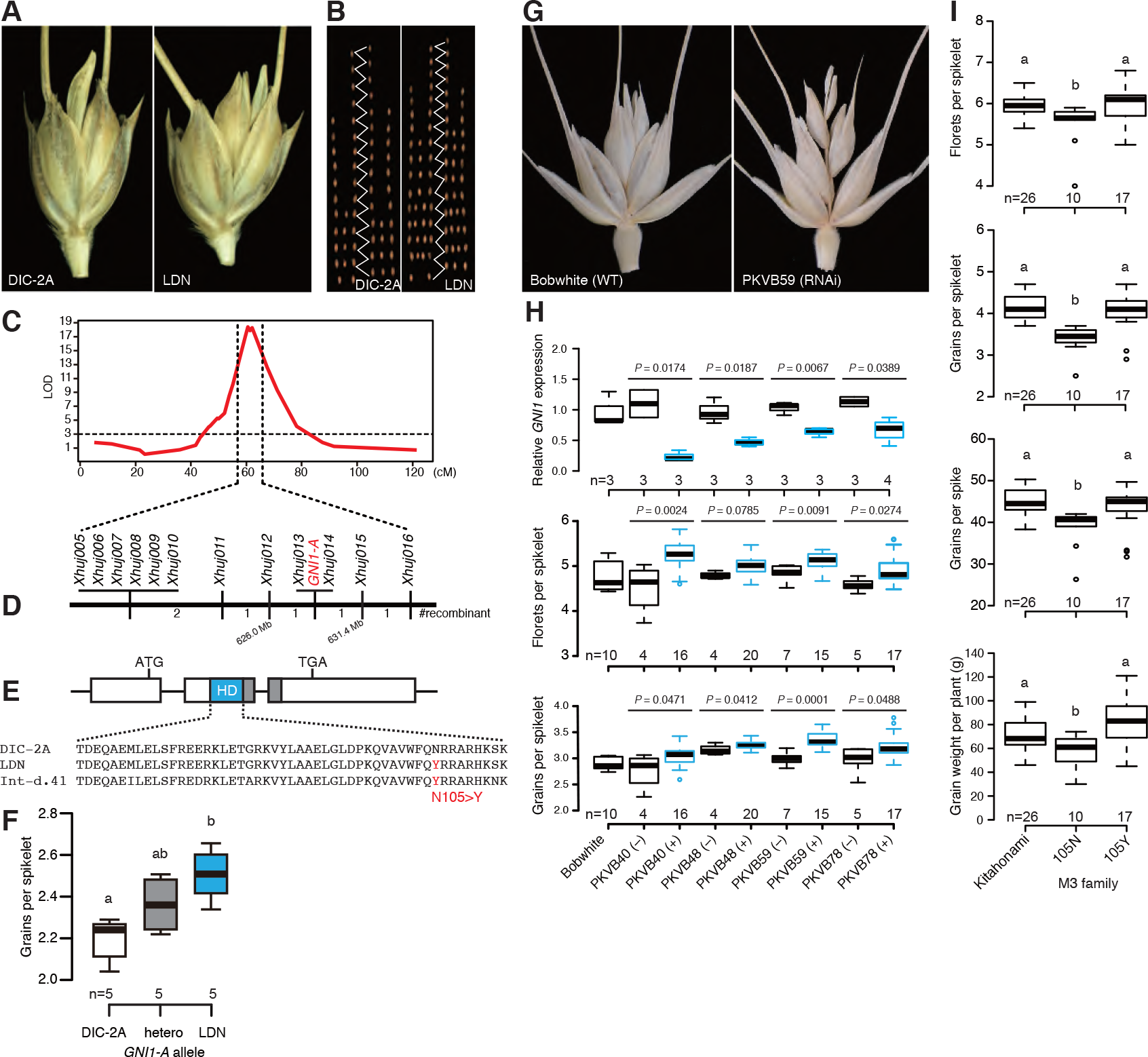
Gene identification responsible for increased grain number per spikelet. (*A*) Representative spikelet of DIC-2A and LDN. (*B*) Number of grains along the spike. (*C*) QTL for grain number per spikelet on chromosome 2AL. (*D*) Fine mapping of the *GNI1-A*. (*E*) Gene structure of *GNI1-A*. The LDN and *Int-d.41* allele carries a single amino acid substitution at the homeodomain (HD). (*F*) Additive effect of the *GNI1-A* alleles in backcrossed recombinants lines. (*G*) Spikelet structure of Bobwhite and transgenic plant with *GNI1-A* RNAi construct. (*H*) Relative *GNI1* expression level (up), number of florets (middle) and grains (bottom) per spikelet of transgenic (+) and non transgenic (−) plants at T1generation. *P*-values are determined by Student’s *t*-tests. (*I*) Phenotypes of TILLING mutants in the M3 family. 105N indicate revertant mutant (functional) allele and 105Y wildtype (reduced-function) allele as Kitahonami. Box edges represent the 25% quantile and 75% quantile with the median values shown by bold lines. Whiskers extend to data less than 1.5 times the interquartile range, and remaining data are indicated by circles. Averages with different letters in (*F*) and (*I*) indicate significant differences as determined by Tukey HSD (*P* ≤ 0.05).

To verify the inhibitory role of *GNI1-A*, we knocked-down its transcript levels using RNA interference (RNAi). An RNAi construct was transformed into hexaploid wheat variety Bobwhite, which contains the 105N allele. We obtained four transformants carrying independent transgenic events and construct-positive plants showed more florets and grains per spikelet on average than construct-negative plants (**Fig. 2G, H**). Reduced transcript levels of *GNI1* were confirmed from all transgenic events (**Fig. 2H**). These results corroborate the hypothesis that functional *GNI1-A* inhibits floret development in wheat. Other traits such as plant height, spike number per plant, spike length, spikelet number per spike and grain size were not significantly changed indicating the spatial specificity of the gene function (***SI Appendix*, Fig. S1**).

### Reduced-function allele of *GNI1-A* improved yield

Kitahonami, one of the highest-yielding bread wheat varieties in Japan, also carried the 105Y allele and showed 4.26 grains per spikelet on average (**Fig. 2I**). Pedigree analysis revealed that the cultivars with the 105Y allele had significantly more grains per spikelet than the other alleles (4.03 for 105Y; 3.21 for 105N and 3.07 for 105K), and the 105Y allele seems to be introduced from UK cultivar, Norman (***SI Appendix*, Fig. S2**). Through TILLING of Kitahonami, we obtained heterozygous M2 plants with an Y105N mutation. As expected, the homozygous 105N mutant plants had significantly fewer florets/grains per spikelet/spike and grain weight per plant compared with carriers of the 105Y allele from the same M3 families (**Fig. 2I**). The change of grain number per spikelet was mainly from the basal and central parts of spike in agreement with the finding in tetraploid wheat. Other traits were not significantly changed (***SI Appendix*, Fig. S3**). These results strongly support the idea that the N105Y mutation of the *GNI1-A* allele contributes to more grains per spikelet due to lowered apical floret abortion during spikelet development.

To reveal the actual effects of the *GNI1-A* allele on grain yield in the field, yield tests were conducted (***SI Appendix*, Fig. S4**). Kitahonami mutants derived from M4 families (105N and 105Y) were compared in two environments (Kitami and Naganuma, Hokkaido, Japan) and 4 and 3 replications in each site, respectively. The results showed that the plants carrying the 105Y allele increased grain plot-yield from 10 to 30% in both sites. Grain number per spike was slightly increased in the 105Y allele and thousand grain weight and number of spikes per plant were not changed (***SI Appendix*, Fig. S4**). Plant biomass was significantly reduced for the 105N allele in the Kitami site. Harvest index was increased in the 105Y allele in both fields. This data indicates that the *GNI1-A* 105Y allele contributes to the improved grain yields under field conditions.

### *GNI1* transcripts accumulate in rachilla

*GNI1* mRNA was localized by *in situ* hybridization of einkorn wheat in spikelet meristems, which generates floret meristems on its lower flanks (**Fig. 3A-F**). After the differentiation of 2^nd^ and 3^rd^ floret primordia at the terminal spikelet stage, *GNI1* signals were localized in the spikelet meristem and apical floret (**Fig. 3B**). *GNI1* signals continued to localize to the spikelet meristem at the white anther stage and were clearly detected in the rachilla bearing the florets (**Fig. 3C-E**). These transcript data imply that GNI1 inhibits apical floret development in spikelets beyond floret 3 at least partly by inhibiting rachilla growth and development.

**Fig. 3.**
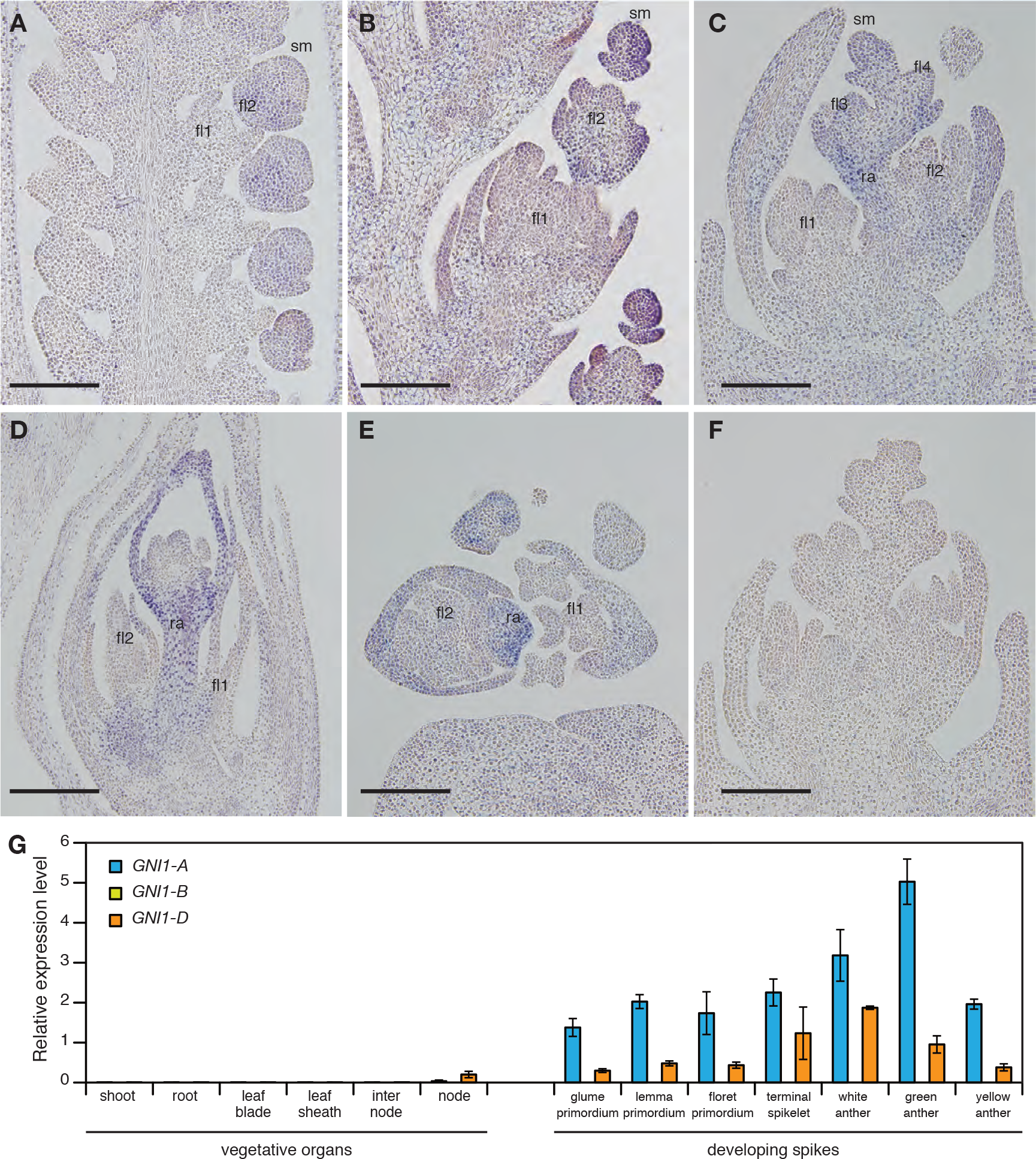
Expression pattern of *GNI1.* (*A*–*F*) *In situ* hybridization of *GNI1* during spike development of einkorn wheat. (*A*) Longitudinal section at floret primordium stage. (*B*) Terminal spikelet stage. (*C*) White anther stage. (*D*) Green anther stage. (*E*) Cross section at white anther stage. (*F*) Control with sense probe. (*G*) Relative expression levels of *GNI1* in bread wheat cv. Bobwhite. Means ± SE of three biological replicates are shown. sm: spikelet meristem, fl: floret, ra: rachilla. Scale bars = 200 μm.

Quantitative RT-PCR analysis showed that *GNI1-A* was predominantly expressed in immature spikes (**Fig. 3G**). Peak mRNA levels were found from the white to green anther stage corresponding to maximum floret primordia number (6). This pattern was conserved among diploid einkorn and tetraploid wheat (***SI Appendix*, Fig. S5**); however, *GNI1-D* showed half of the *GNI1-A* transcript levels and *GNI1-B* transcript levels were negligible in the organs examined although its gene structure is conserved (***SI Appendix*, Fig. S6**).

To better understand the underlying molecular mechanism related to the *GNI1-A* mutation, an RNA-seq profiles of the 105N and 105Y alleles derived from Kitahonami-mutant M4 families were compared. The results revealed that nitrogen metabolic processes, sucrose metabolic processes, and G-protein beta/gamma-subunit complex binding related genes were up-regulated in the 105Y allele compared to 105N mutant allele supporting the increased grain number phenotype in 105Y (**Datasets S1 and S2**). In accordance with the improved floret fertility found in 105Y, we found that the florigenic *Flowering locus T* homolog, *FT-D1*, was upregulated; while both other homoeo-alleles, *FT-A1* and *FT-B1*, were expressed at the same level in spikes at white and green anther stages. Very similar expression patterns were observed in public RNA-seq database (***SI Appendix*, Fig. S7**). These results suggest that *FT1* may function as a floral promoting factor during early floret development in wheat; while differential upregulation of *FT-D1* might be the consequence of higher floral activity in a *GNI1-A* 105Y allele background.

### Natural variation of *GNI1-A* in wheat

To interrogate natural variation at the *GNI1-A* locus, 72 tetraploid wheat accessions including wild and domesticated emmer and durum wheat were sequenced. The panel revealed nine haplotypes (**Fig. 4A**, ***SI Appendix*, Table S2**). Only durum wheat carried the 105Y allele and showed a significantly higher number of grains per spikelet compared with wild and domesticated emmer wheat (**Fig. 4B**). Furthermore, we examined grains per spikelet in three different growth conditions. The results indicated that lines carrying the 105Y allele had a higher number of grains per spikelet in all environments with high broad-sense heritability (*H*_*bs*_ = 0.8), and stable trait expression even under low-yielding conditions (Ruhama site; **Fig. 4C**). In bread wheat, sequencing revealed three haplotypes among 210 European winter wheat cultivars. Hap1 and 2 contained the 105N allele and Hap3 contained the 105Y allele (**Fig. 4D**, ***SI Appendix*, Table S3**). Number of florets at the green anther stage, indicative of maximal grain number potential, was not significantly different between these three haplotypes (**Fig. 4E**). Cultivars carrying Hap3 showed more grains per spikelet at apical (2.97) and central (3.95)positions of the spike and on average (3.52) compared to the other cultivars carrying Hap1 (2.45 at apical, 3.58 at central, and 3.18 on average) and Hap2 (2.53 at apical, 3.64 at central, and 3.22 on average)(**Fig. 4F**). Furthermore, Hap3 cultivars showed more grains per spike, a higher spike fertility index and ratio between spike and stem dry weight due to reduced stem and leaf dry weight (***SI Appendix*, Fig. S8**).

**Fig. 4.**
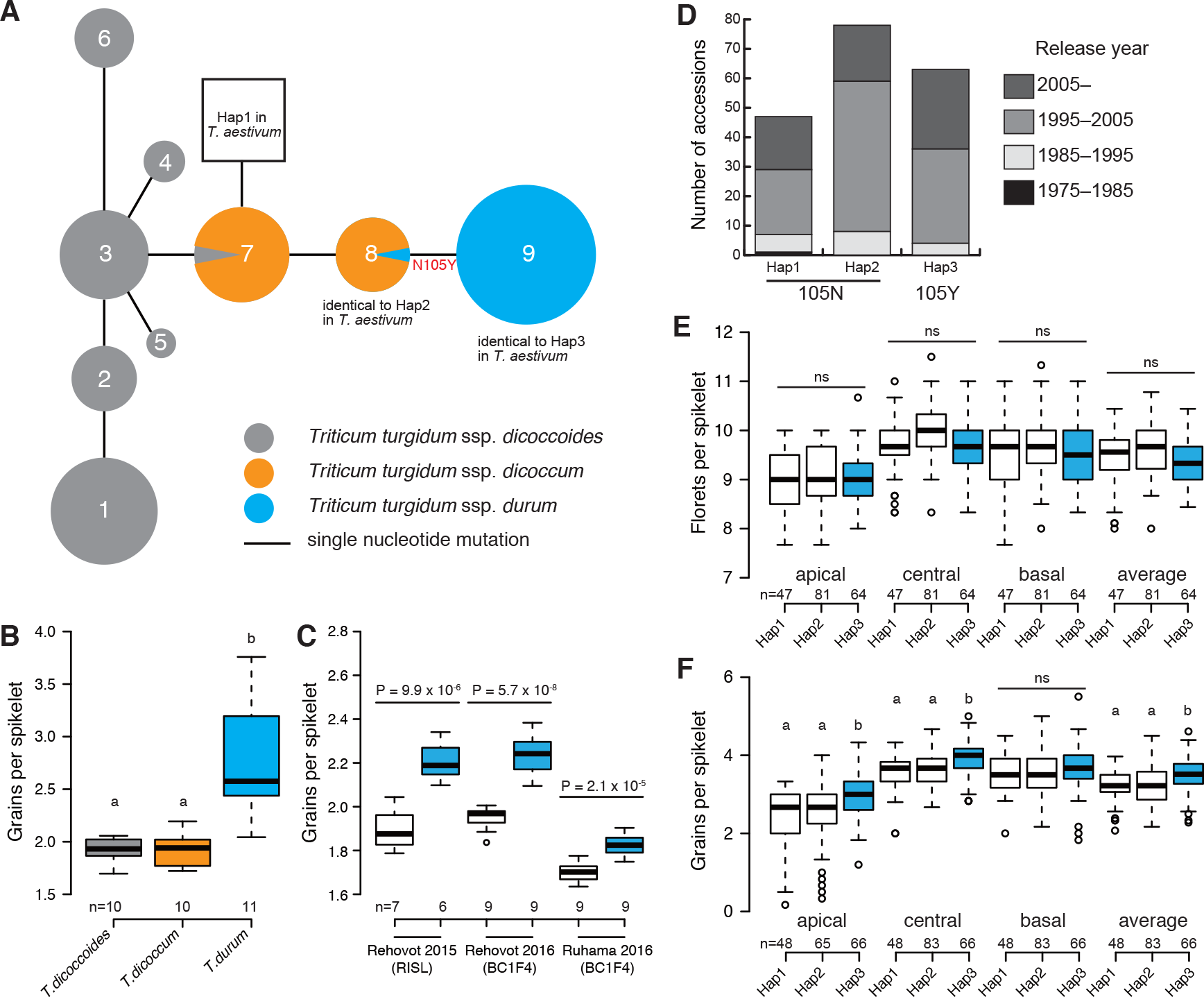
Natural variation of *GNI1-A*. (*A*) Haplotype network analysis of *GNI1-A* in tetraploid wheat. The median-joining network was constructed from the haplotypes based on the resequencing of 35 wild emmer, 14 domesticated emmer and 23 durum wheat accessions. Circle sizes correspond to the frequency of individual haplotypes. Lines represent genetic distances between haplotypes. (*B*) Grains per spikelet in tetraploid wheat. (*C*) Effect of *GNI1-A* allele in tetraploid wheat across environments. 13 RISLs and 18 BC1F4 lines were used. *P*-values are determined by Student’s *t*-tests. (*D*) Frequency of three haplotypes in 210 EU winter bread wheat cultivars. (*E*) Number of florets per spikelet at green anther stage. (*F*) Hap3 (105Y) cultivars showed more grains per spikelet. Different letters in (*B*), (*E*) and (*F*) indicate significant differences as determined by Tukey HSD (*P* ≤ 0.05). ns; not significant.

### Evolution of *GNI1* in Triticeae

To shed more light on the evolution of *GNI1* (i.e. *HOX1*) in Triticeae, diverse collections of wild species were sequenced by nucleotide sequence capture. All 14 genera examined in this study had *HOX2* homologs, which is in line with the fact that *HOX2* is highly conserved in the orthologous region in grass species (22). *HOX1* homologs were only present in seven genera; *Hordeum*, *Dasypyrum*, *Secale, Taeniatherum*, *Aegilops*, *Amblyopyrum*, and *Triticum* (***SI Appendix*, Fig. S9**). There were no taxa among Triticeae carrying only *HOX1*. These results indicate that the duplication of the ancestral gene may have occurred after the divergence of Triticeae. There might be two gene lineages: one including seven genera, such as *Psathyrostachys*, *Pseudoroegneria*, *Agropyron*, maintained one copy without duplication event, while the other seven genera, such as *Hordeum*, *Secale*, *Triticum*, retained two copies.

To query the relationship between the number of fertile florets and ploidy level, we investigated diploid, tetraploid and hexaploid wheat relatives. As expected, a higher ploidy level increased floret fertility except for *Aegilops* species. Diploid A-genome progenitor, *T. urartu* produced three florets and a single grain per spikelet (**Fig. 5A, B**), while *Aegilops* species (S-, D-genomes) showed three to six florets and one to four grains per spikelet (**Fig. 5A, B**). Tetraploid wheat showed four to five florets and two to three grains. Hexaploid wheat showed five to seven florets and three to five grains (**Fig. 5A, B**). To further test the relationship between fertile floret number and *GNI1* transcript levels, qRT-PCR was performed. We found a negative trend between the number of florets or grains per spikelet and *GNI1* mRNA expression level (**Fig. 5C**), suggesting that the lower *GNI1* expression results in higher number of fertile florets in both diploid and polyploid wheat. The implication was a functional diversification not only between *HOX1* and *HOX2* but also among *HOX1* genes in the *HOX1*-carriers in the process of their speciation.

**Fig. 5.**
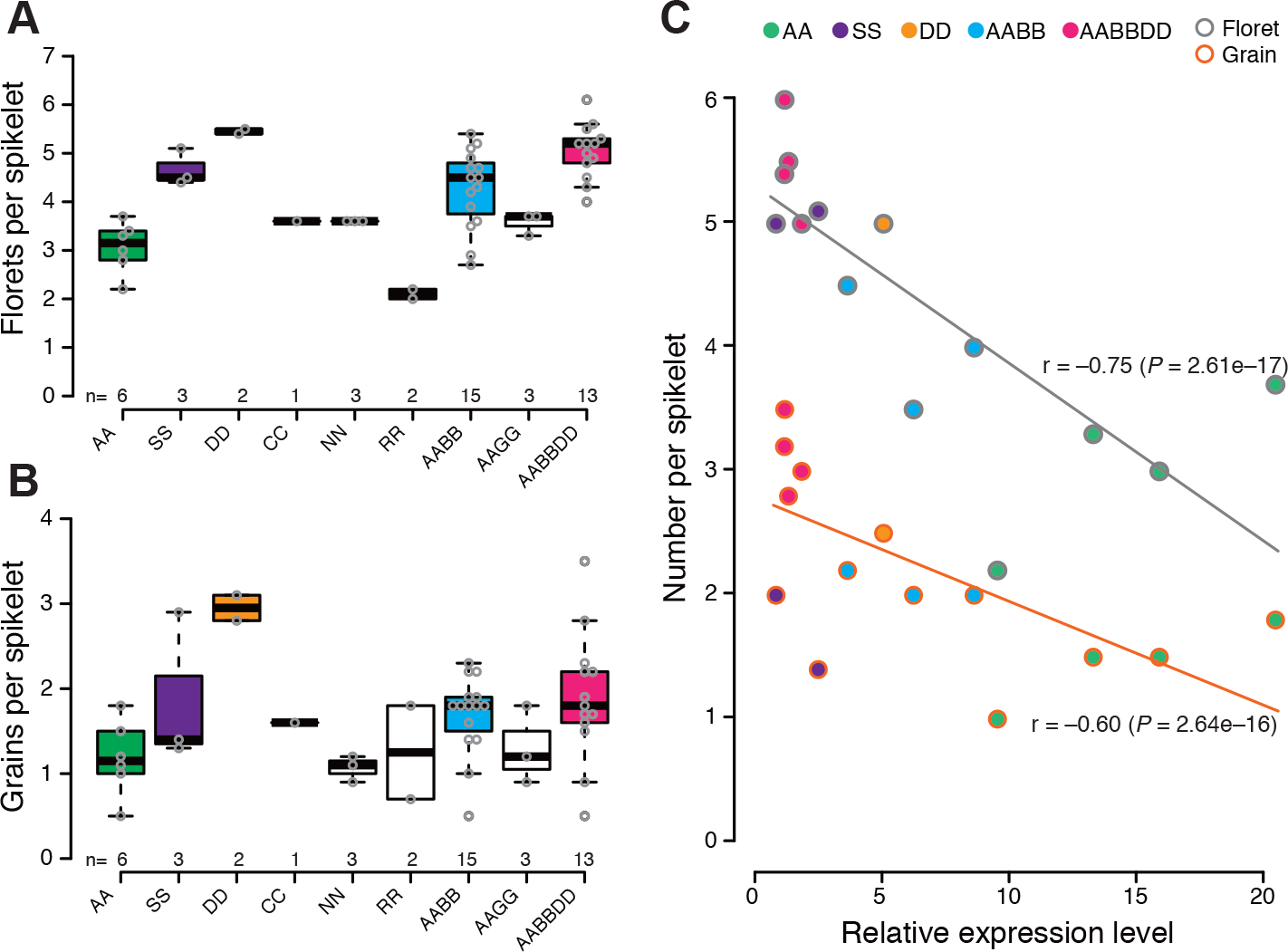
Functions of *GNI1* in diploid, tetraploid and hexaploid species. (*A*) Floret number per spikelet. (*B*) Grain number per spikelet. (*C*) Relationship between floret/grain number per spikelet and expression level of *GNI1*.

## Discussion

In this study, we show that the *GNI1* gene is a key determinant of grain number by controlling floret fertility in wheat. *GNI1* encodes an HD-Zip I transcription factor that is preferentially expressed during rachilla growth and development. Reduced-function mutation of *GNI1* increases the grain number and yield without adverse effects on the number of spikes, number of spikelet per spike and grain size. These results strongly suggest that improving floret fertility (i.e. relieving floret abortion) is a promising breeding strategy for enhancing grain yield in the unbranched “spike” type inflorescence like wheat. Increased floret numbers were observed in mutated *APETALA2* (*AP2*) gene family transcription factors, such as *ids1* in maize and *Q* in wheat, which control the spikelet meristem fate; however, their floret fertility is relatively low (23, 24). This indicates that determining appropriate number of floret primordia, not indeterminate number is useful for improving grain number. Optimization of both HD-Zip I (i.e. *GNI1*) and *AP2* genes may help improving floret fertility, but its relationships remain elusive. Identifying network genes under the control of *GNI1* and investigating the other genes responsible for increased floret fertility will be important to further enhance grain yield in wheat.

HD-Zip class I genes, including *GNI1* in wheat, have evolved through a series of gene duplications, functionalizations and mutations (25–27). *GNI1* evolved through gene duplication in Triticeae after the separation of Triticeae and Bromeae. Only seven genera, such as *Hordeum*, *Secale*, *Triticum,* out of fourteen genera tested in this study experienced the gene duplication. Seven genera, such as *Psathyrostachys*, *Pseudoroegneria*, *Agropyron*, maintained one copy without duplication event. This notion was well supported by plenty of repetitions (species and accessions) from each genus. Truncated HD-Zip I gene sequences were not evident from the exome sequencing, indicating that the missing of the HD-Zip I in the second group was not a consequence of spontaneous, independent deletions of the genes, but rather they were not duplicated from the origin of their common ancestor. Interestingly, only plants of the first group (with duplicated copies) were domesticated as cereal crops.

*GNI1* is the ortholog of the barley *Vrs1* gene, which controls lateral floret fertility. *Vrs1* inhibits the development of lateral florets, especially pistil (21, 27), whereas loss-of-function mutations produce up to three times as many grains per rachis node. While *Vrs1* mRNA signals accumulate in rachilla and pistil (27), *GNI1* mRNA is highly accumulating during rachilla growth and development, suggesting that this is its ancestral role. In addition, barley *Vrs1* may have acquired a specific transcriptional expression pattern characteristic for the evolution of lateral spikelet/floret/pistil development within the genus *Hordeum*. Our expression analysis further suggests that *Aegilops* species (S and D genome) are likely to be pseudogenized, whereas diploid *Triticum* (A genome) and *Hordeum* (H genome) species are functionalized (27). These *GNI1/Vrs1*-mediated floral changes follow a theory of regulatory evolution: that morphological diversity is driven by changes in gene expression pattern that minimize pleiotropic effects while simultaneously maximizing adaptation (28, 29).

The single amino acid substitution in a conserved domain causing the reduced-function mutation (105Y) of *GNI1-A* was selected after the divergence of durum wheat. Previous study showed that six-rowed barley is originated from domesticated two-rowed type through mutations of *Vrs1* (21). Taken together, these data suggest that the mutation for increased grain number have undergone parallel selection during post-domestication of wheat and barley (30). Its high allele frequency among durum wheats (96%) supports the notion for a selection towards increased grain number. The mutant allele (105Y) is absent in any wild and domesticated emmer wheat analyzed so far (among 64 accessions) suggesting that the mutation had negative effect on plant’s fitness under natural conditions. During wheat evolution under domestication, the mutant allele become more beneficial in human-made environment, i.e. increased yields, and such strong selection pressure resulted in fixation of the mutant allele in durum wheat. The wildtype (105N) and mutant (105Y) alleles occurred in a 2:1 ratio in the European winter bread wheat gene pool, suggesting that the 105Y allele may have entered the breeding programs more recently. *GNI1* heritability was high and its allele effect was stable across environments (**Fig. 4C** **and *SI Appendix*, Fig. S4**), supporting the usefulness of the reduced-function allele in wheat breeding programs to increase grain yield around the world.

## Methods

### QTL mapping

A population of 94 Recombinant Inbred Substitution Lines (RISLs) derived from a cross between durum wheat [*T. turgidum* ssp. *durum* (Desf.) MacKey] cv. Langdon (LDN) and the chromosome substitution line DIC-2A, which contains the chromosome 2A of wild emmer wheat [*T. turgidum* ssp. *dicoccoides* (Körn.) Thell.] accession FA-15-3 (ISR-A) against the LDN genetic background (20), was used for QTL analysis. The experiment was conducted in an insect-proof screen-house at the experimental farm of the Hebrew University of Jerusalem in Rehovot, Israel (34°47′ N, 31°54′ E; 54 m above sea level). A complete random block design, with three replicates (single plant) was applied. Three spikes per plant were characterized for number of spikelets and number of grains per spike. Number of grains per spikelet was calculated accordingly. Linkage analyses and map construction were performed based on the evolutionary strategy algorithm included in the MultiPoint package (http://www.MultiQTL.com), as previously described (31). QTL analysis was performed with the MultiQTL package, using the general interval mapping for RIL-selfing populations. The hypothesis that one locus on the chromosome has an effect on a given trait (H_1_) was compared with the null hypothesis (H_0_) that the locus had no effect on that trait. Once the genetic model was chosen, 10,000 bootstrap samples were run to estimate the standard deviation of the main parameters: locus effect, its chromosomal position, its LOD score and the proportion of explained variation (PEV).

### Fine mapping

For fine mapping of *GNI1-A* locus, three RISLs containing the DIC-2A allele in the QTL interval (RISL #4, #63, #102) were backcrossed to LDN and then self-pollinated to produce BC_1_F_2_. To identify recombinants within the genomic region including the *GNI-1A* locus, 1006 BC_1_F_2_ plants were screened with microsatellites markers *Xgwm558* and *Xcfa2043*. In the next generation, BC_1_F_3_ families (8 seedlings per family) were screened again with markers *Xgwm558*, *Xcfa2043* and *Xhbg494*. Homozygous recombinants were genotyped with 12 additional markers (***SI Appendix*, Table S4**) developed based on polymorphism between wild and domesticated emmer wheat genomes (32). The phenotype of 17 BC_1_F_4_ homozygous recombinants was scored in the field. A split plot, complete random block design (*n*=5), was carried out in insect-proof screen-house at the experimental farm of the Hebrew University of Jerusalem in Rehovot, Israel. Each plot (1×1 meter) contained twenty plants per line. Five spikes per plot were characterized for number of spikelets per spike and number of grains per spike. Number of grains per spikelet was calculated accordingly.

To examine the effect of the *GNI1-A* allele across environments, three field experiments were analyzed. The first experiment was conducted in 2015-16 in Rehovot, with 13 RISLs and their parental lines, LDN and DIC-2A. Two field experiments were conducted in 2016-17, in Rehovot and in Ruhama (31°29’ N, 34°42’ E, 166m above sea level) using 18 BC_1_F_4_ lines and LDN. All experiments were conducted as described above.

### Transformation of wheat

We used a 323-bp fragment from the *GNI1*-*A* as RNA interference (RNAi) trigger. This fragment was amplified using specific primers (***SI Appendix*, Table S4**) and cloned it into a pENTR D-TOPO vector (Thermo Fisher Scientific, Waltham, MA USA). The fragment was transferred by LR recombination reaction into a pANDA-b vector in which transgene expression was under control of the maize *Ubiquitin1* promoter (33). Transgenic wheat plants were produced by a particle bombardment method using immature embryos of spring-type bread wheat variety Bobwhite S-26 obtained from CIMMYT (34). Transgene-positive plants were confirmed by PCR using vector specific primers (***SI Appendix*, Table S4**).

### TILLING

Winter-type bread wheat cv. Kitahonami was used as donor plants for mutagenesis. The mutations were induced by gamma-ray irradiation (250 Gy) using mature grains. M1 plants were grown in greenhouse and all M2 grains were harvested. The DNA was extracted from M2 plants and used as template of TILLING. *GNI1-A* specific primers (***SI Appendix*, Table S4**) were used to amplify the fragment. M3 family was evaluated in the NIAS experimental field in Tsukuba, Japan (36°01’ N, 140°06’ E).

### Yield trials

Kitahonami mutants (105Y allele and 105N allele) at M4 generation were used. The plants were grown in the Kitami Agricultural Experiment Station (43°44’N, 143°43’W, Kitami, Hokkaido, Japan) and the Central Agricultural Experiment Station (43°3’N, 141°45’W, Naganuma, Hokkaido, Japan). At Kitami, each genotype consisted of four plots (5.4m^2^ per plot, 200 grains per m^2^). At Naganuma, each genotype consisted of three plots (4.8m^2^ per plot, 200 grains per m^2^). 156 kg N, 175 kg P, 70 kg K/ha at Kitami and 140 kg N, 125 kg P, 50 kg K/ha at Naganuma were applied.

### RNA extraction and Absolute Quantitative Real-Time PCR

Immature spikes were developmentally staged by observation under the stereoscopic microscope (35). Total RNA was extracted using TRIzol (Invitrogen) and quantified using NanoDrop 1000 (Thermo Fisher Scientific). To remove genomic DNA contamination, RNA was treated with RNase-free DNase (Takara Bio). First-strand cDNA was synthesized with SuperScript III (Invitrogen) and first-strand cDNA derived from 25 ng RNA was used as template. Transcript levels of each gene were measured by quantitative real-time PCR using a StepOne Real-Time PCR System (Applied Biosystems) and THUNDERBIRD SYBR qPCR Mix Kit (Toyobo) according to the manufacturers’ protocols. Primers used for qRT-PCR are listed in ***SI Appendix*, Table S4**. Each gene fragment was cloned into pBluescript II KS (+). Plasmid DNA harboring each gene fragment was used to generate standard curves for absolute quantification. qRT-PCR analysis was performed at least three times for each sample. Biological replicates of at least three independent RNA extractions per sample were performed. The *Actin* gene was used to normalize the RNA level for each sample.

### *In situ* mRNA hybridization analysis

The *GNI1* gene segment comprising part of the 3′-UTR (300 bp) was amplified from cDNA isolated from immature wheat spikes using specific primers (***SI Appendix*, Table S4**). The PCR product was cloned into the pBluescript II KS (+) vector (Stratagene). Two clones with different insert orientations were linearized with *Eco*RI and were used as templates to generate antisense and sense probes using T3 RNA polymerase. *In situ* hybridization was conducted as in Komatsuda, *et al.* (21).

### RNA-sequencing and analysis

For RNA-sequencing (RNA-seq), immature inflorescences at the white anther stage and green anther stage of Kitahonami M4 families (105Y allele and 105N allele) were collected. Total RNA was extracted using TRIzol (Invitrogen) and treated with DNase I (Roche). Quality of RNA was measured using an Agilent 2100 Bioanalyzer (Agilent Technologies). RNA-seq libraries were prepared using the NEBNext UltraTM RNA Library Prep Kit for Illumina (NEB, USA) and sequenced using a HiSeq 4000 (Illumina).

Transcript abundance (transcripts per million reads) was quantified by pseudo-alignment against the representative transcripts of high-confidence and low-confidence genes annotated on the IWGSC RefSeq V1 pseudomolecules (36) using Kallisto version 0.43.1 (37). Abundance estimates were imported into the R statistical environment (38). Analysis of differential gene expression was carried out with Limma-voom (39, 40). Differentially expressed genes (DEGs) were required to have log2 fold change ≥ 2 or ≤ −2 between contrasted conditions and an adjusted *P*-value ≤ 0.05 after Benjamini-Hochberg correction. Enrichment of gene ontology (GO) terms was analyzed with topGO (41) using the GO term assignment of IWGSC, 2018.

### Haplotype analysis

For resequencing, 72 tetraploid wheat accessions including 35 wild emmer, 14 domesticated emmer and 23 durum were used (***SI Appendix*, Table S2**). Out of 72, 30 accessions were evaluated for the grain number per spikelet. A haplotype analysis was performed based on resequencing data of the *GNI1-A* (1,053 bp) from tetraploid wheat. Sequence alignments were performed with ClustalW using MEGA7 software (42). A median-joining network (43) was constructed using the software programs DNA Alignment version 1.3.3.2 and Network version 5.0.0.1 (Fluxus Technology) with default parameters (epsilon = 0; frequency .1 criterion = inactive; The ratio of transversion:transition = 1:1; Criterion = Connection cost; External rooting = inactive; MJ square option = inactive). A total of 210 European winter bread wheat cultivars (***SI Appendix*, Table S3)**were used for resequencing *GNI1-A* with specific primers (***SI Appendix*, Table S4**). Phenotypic data from Guo, *et al.* (19) was reanalyzed with *GNI1-A* haplotype data (**Dataset S3**).

### Phylogenetic analysis

For phylogenetic analysis, 139 individuals of 35 species and 14 genera of Triticeae and *Brachypodium* and *Bromus* as outgroup taxa were used (***SI Appendix*, Table S5**). The sequence capture was performed as previously described(44). For each taxon, a set of overlapping probes was designed to cover each sequence at least four times. Probe sequences are listed in **Dataset S4**.

The sequence assembly was performed using GENEIOUS v. 10.0.9 (45). For the diploid and autotetraploid individuals, the reads were mapped simultaneously to barley *Vrs1* (AB478778) and *HvHox2* (AB490233) using the “Medium-Low Sensitivity” parameter. Only reads for which both sequences of a pair mapped were kept. For heterozygous diploid and allotetraploid individuals, the haplotype phasing consisted of mapping followed by *de novo* assembly to obtain two alleles or homoeologues per locus as previously described (46) with the following modifications: diploid heterozygous individuals were phased individual-wise. As a control the two obtained alleles from each individual were then used as references for mapping the individual’s reads. For allotetraploid individuals, the phasing was performed only on the individual of a species with the highest sequence coverage. The retrieved homoeologues were then used as references (KU-8939, PI 428093). To control for potential chimeras the resulting assemblies were carefully inspected and their contigs were aligned with MAFFT v. 7.308 (47). For *Triticum aestivum*, *GNI1* and *TaHox2* from Chinese Spring were used as references for read mapping. All assemblies were inspected for misalignments and coverage inconsistencies. Consensus sequences were called using the ‘Highest Quality’ threshold in GENEIOUS.

The multiple sequence alignment was performed in GENEIOUS v. 10.2.3 using MAFFT. The alignment was checked by eye and trimmed to the coding sequences. The best-fit model of nucleotide substitution was selected using JMODELTEST2 (48). The Bayesian information criterion (49) selected K80 + G out of 24 tested models. The Bayesian phylogenetic inferences was performed with MrBayes version 3.2.6 (50) on CIPRES (Cyberinfrastructure for Phylogenetic Research) Science Gateway 3.3 (51, 52). Two parallel Metropolis coupled Monte Carlo Markov chain analyses with four chains were run for six million generations sampling trees every 500 generations. Convergence of the runs was assessed using the standard deviation of split frequencies being <0.01. The continuous parameter values sampled from the chains were checked for mixing using Tracer v1.6 (http://beast.bio.ed.ac.uk/Tracer). A consensus tree was computed in MrBayes after removal (burn-in) of the first 25% of trees. The consensus tree was visualized in FIGTREE v. 1.4.2 (http://tree.bio.ed.ac.uk/software/figtree) using the node representing the most recent common ancestor of *Brachypodium* and the Triticeae as the root.

### Accession numbers

The gene sequences are available from DNA Data Bank of Japan (DDBJ) under accession numbers AB711370 – AB711394, AB711888 – AB711913 and NCBI GenBank under accession numbers MH134165 – MH134483. The RNA-seq data have been submitted to the European Nucleotide Archive under accession number PRJEB25119.

## Acknowledgments

Grains of the RISL population were kindly provided by Steven S. Xu and Justin D. Faris (USDA-ARS, Fargo, ND USA). DNA samples and grains of the Kitahonami TILLING population were kindly provided by Yoko Ono, Fuminori Kobayashi and Hirokazu Handa (NARO). The 210 EU winter wheats were kindly provided by Marion S. Röder (IPK). We thank Hiroyuki Tsuji (Yokohama City University) for providing pANDA-b vector. We also thank NBRP-wheat, Kyoto, Japan; the Nordic Genetic Resources Center (NordGen), Alnarp, Sweden; and the IPK Genebank, Gatersleben, Germany; and USDA, Idaho, USA for providing germplasm. We thank Shinji Kikuchi (Chiba University), Harumi Koyama, Mari Sakuma (NIAS), Corinna Trautewig and Anne Fiebig (IPK) for excellent technical support. This work was supported by grants from the Genomics for Agricultural Innovation Program of the Ministry of Agriculture, Forestry, and Fisheries of Japan (TRS1002 to S.S. and T.K.) and a Grant-in-Aid from the Japan Society for the Promotion of Science (JSPS) Postdoctoral Fellow for Research Abroad (to S.S.); a Grant-in-Aid for Young Scientists (B) (no. 16K18635 to S.S.); the Chief Scientist of the Israel Ministry of Agriculture and Rural Development 20-10-0066 to Z.P.), the U.S. Agency for International Development Middle East Research and Cooperation (M34-037 to Z.P.) and a grant from the German Research Foundation (BL462/10).

## Author contributions

S.S., G.G., Z.P., T.S. and T.K. designed research. S.S., G.G., Z.G., T.O., A.T., K.S., N.B., J.B., G.H., S.O., H.J., Y.Y., I.A., Z.P. performed experiments. M.M. analyzed NGS data. S.S., T.S. and T.K. wrote the manuscript with contributions from all coauthors.

